# A coarse-grained approach to model the dynamics of the actomyosin cortex

**DOI:** 10.1101/2021.05.20.444937

**Authors:** Miguel Hernández-del-Valle, Andrea Valencia-Expósito, Antonio López-Izquierdo, Pau Casanova-Ferrer, Pedro Tarazona, Maria D. Martín-Bermudo, David G. Míguez

## Abstract

The dynamics of the actomyosin machinery is at the core of many important biological processes. Several relevant cellular responses such as the rhythmic compression of the cell cortex are governed, at a mesoscopic level, by the nonlinear interaction between actin monomers, actin crosslinkers and myosin motors. Coarse grained models are an optimal tool to study actomyosin systems, since they can include processes that occur at long time and space scales, while maintaining the most relevant features of the molecular interactions. Here, we present a coarse grained model of a two-dimensional actomyosin cortex, adjacent to a three-dimensional cytoplasm. Our simplified model incorporates only well characterized interactions between actin monomers, actin crosslinkers and myosin, and it is able to reproduce many of the most important aspects of actin filament and actomyosin network formation, such as dynamics of polymerization and depolymerization, treadmilling, network formation and the autonomous oscillatory dynamics of actomyosin. Furthermore, the model can be used to predict the *in vivo* response of actomyosin networks to changes in key parameters of the system, such as alterations in the anchor of actin filaments to the cell cortex.

## Introduction

The interactions that govern the dynamics of biological systems take place at size and time scales that can differ several orders of magnitude. At the macroscopic level, computational characterization and analysis of biological processes relies on stochastic or deterministic differential equations. At the atomic resolution level, Molecular Dynamics (MD) numerical simulations are the main tool, but are restricted to very few molecules and short time scales. In between both scales, models that involve many molecules and/or larger length (typically larger than 10 nm) and time scales (longer than 100 ns), are often based on coarse-grained (CG) approximations (Wu and Shea, 2011).

CG models can be defined as a mathematical representation of a system made of simplified versions of its more relevant sub-components and interactions (Ayton *et al*, 2007). These type of models constitute an optimal tool to study system properties that arise from the molecular aspects of the interacting components (Tozzini, 2005), such as aggregation, polymerization or self-assembly (Murtola *et al*, 2009). In the context of the cell cytoskeleton, (Kmiecik *et al*, 2016) microtubule formation (Gebremichael *et al*, 2008), microtubule mechanics (Ding and Xu, 2011; Theisen *et al*, 2013), dynamics (Ji and Feng, 2011), and interaction with kinesin motors (Hyeon and Onuchic, 2011) have been addressed computationally using a CG approximation.

The other main component of the cytoskeleton, the actomyosin machinery, has been extensively studied using CG models, such as the assembly (Chu and Voth, 2006) and allostery (Chu and Voth, 2005) of actin filaments (F-actin) from globular actin (G-actin), its reorganization in networks and bundles in the cell cortex (Shin *et al*, 2009), the fluidization of the actin cytoskeleton (Li *et al*, 2007) as well as other responses under stretching (Kang *et al*, 2011). In addition, Vavylonis et al. propose a numerical model for actomyosin contraction where some of the features of actomyosin network formation are modeled using a Monte Carlo approximation (Vavylonis *et al*, 2008).

The effects of crosslinker proteins (Fallqvist *et al*, 2014; Kim *et al*, 2009), myosin V and Myosin II (Taylor and Katsimitsoulia, 2010) have also been studied using a CG approximation (Ennomani *et al*, 2016).

One of the most interesting properties of the actomyosin machinery is their ability to undergo periodic oscillations in their concentration (Solon *et al*, 2009; Blanchard *et al*, 2010; He *et al*, 2010; Valencia-Expósito *et al*, 2016). These oscillations take place as pulsating reorganization of twodimensional networks of actin and myosin in the cell cortex, a specialized layer attached to the inner plasma membrane that plays a central role in cell motility and cell shape control.

This type of autonomous oscillatory dynamics is ubiquitous in biology at many levels, from gene expression (Ciliberto *et al*, 2005; Swinburne *et al*, 2008) to circadian clocks (Nagoshi *et al*, 2004) and nervous impulses (Marder and Bucher, 2001). At the cellular level, spontaneous mechanical oscillations have been reported in a variety of cell types, including cardiomyocytes (Fabiato and Fabiato, 1978; Viatchenko-Karpinski *et al*, 1999), fibrils of ordinary skeletal muscles (Yasuda *et al*, 1996; Anazawa *et al*, 1992; Ishiwata *et al*, 2007), insect muscle cells (Jewell and Ruegg, 1966) and eukaryote tissue culture cells (Sedzinski *et al*, 2011). In the context of animal development, autonomous oscillations have been shown to govern the dynamics of the segmentation clock during somitogenesis (Jiang *et al*, 2000; Kaern *et al*, 2004), the neurogenesis in the developing cortex (Banerjee and Ellender, 2009) or the overall size of the embryo (Lauschke *et al*, 2013). In addition, the spindle oscillations during asymmetric division in C. Elegans (Pecreaux *et al*, 2006) arise from periodic mechanical coordination of force-generating motors (Grill *et al*, 2005). Apical constriction (Martin *et al*, 2009) in ventral furrow cells pulsates during *Drosophila* mesoderm invagination (Martin *et al*, 2009). Periodic contraction and expansion of amnioserossa cells drives tissue movement during dorsal closure (Solon *et al*, 2009; Blanchard *et al*, 2010). In addition, elongation of the *Drosophila* egg chamber requires periodic oscillations of the actomyosin network present on the basal side of follicle cells (He *et al*, 2010; Valencia-Expósito *et al*, 2016).

The basic machinery for these mechanical oscillations relies on the interactions that govern the dynamics of the actomyosin cell cortex. It has been shown that cooperative myosin motors acting over an actin filament oscillate with a frequency of few tenths of Hertzs and around 10 nm of amplitude (Jülicher and Prost, 1997; Sato *et al*, 2011), but the mechanism underlying the much slower and larger actomyosin oscillations responsible for cell shape oscillations is still poorly understood (Solon *et al*, 2009; Blanchard *et al*, 2010). Recently, several theoretical studies have proposed a combination of mechanical and biochemical interactions for emergence of periodic constriction of the apical cell cortex of amnioserosa cells during dorsal closure in *Drosophila*. An initial model proposes the interplay between myosin and an unknown signaling molecule that oscillates due to the interaction with myosin combined with coupling between neighboring cells (Solon *et al*, 2009). Recently, several contributions proposed a mechano-chemical cell autonomous mechanism based on previous models (Sedzinski *et al*, 2011) of pulsating dynamics acting in dividing cells, where coupling between actin turnover and cell area is assumed (Machado *et al*, 2014). Other recent approaches reproduce actomyosin pulsations combining the effect of nonlinear actin and myosin turnover, an elastic restoring force and a viscous damper (Dierkes *et al*, 2014). Other mechanochemical approaches specific to the oscillations in the basal cell cortex in *Drosophila* follicle cells in the developing egg chamber assume cells as springs acting against a potential internal pressure of the egg chamber (Koride *et al*, 2014).

We have recently proposed an alternative model for actomyosin oscillations based on aggregation and dissociation of actin filaments mediated by the mechanical action of myosin. The model reproduces the spontaneous oscillations in actomyosin concentration observed on the basal side of *Drosophila* follicle cells based simply on F-actin aggregation and myosin induced disassembly of the network (Valencia-Expósito *et al*, 2016). In addition, this simplified model reproduces the experimental features of several mutant conditions, such as changes in myosin activity (Valencia-Expósito *et al*, 2016), and in integrin levels (Santa-Cruz Mateos *et al*, 2020).

In this paper, we use a numerical Monte-Carlo approximation to develop a CG model of the actomyosin cortex, based on three main interactions: the polymerization and depolymerization of F-actin from G-actin, the assembly of networks of F-actin attached to the inner cellular membrane mediated by actin binding proteins, and the disassembly of F-actin networks mediated by myosin. Our model successfully reproduces key aspects of the actomyosin cortex, such as polymerization dynamics, filament size distribution, treadmilling, cooperative F-actin assembly and network formation above a threshold concentration. The model also reproduces the periodic autonomous assembly of the actomyosin cortex observed in many biological systems, as well as the effect in the oscillations due to changes in the levels of key proteins, such as Myosin and Integrins.

Despite the highly complex biological regulation of the actomyosin machinery, the model shows that many of the most relevant aspects of the actomyosin cortex can be explained based on simplified coarse-grained interactions between actin, myosin and crosslinker molecules.

## Results

### The model reproduces the dynamics of F-actin polymerization

In this section, we use a first version of the framework with G-Actin as the only ingredient to study the dynamics of F-Actin formation, its dependence on key model parameters, and how it compares with well established experimental observations. Details of how the model is implemented are explained in the Methods section, and summarized in Box 1.

#### Box 1

##### F-Actin polymerization

(a) A position in the grid [*x, y*] is sampled at random. (b) If the position is empty, the probability of incorporation of a G-Actin from the cytoplasm to this position in the grid is computed as *P*^+^ = *e*^*μ*_1_^; (c) If the incorporation is successful, the orientation of the molecule in the grid is chosen at random between four possible configurations: north, south, east or west; (d) When a G-actin molecule enters the grid in front of a F-Actin with the same orientation, a bond (with energy *E*_0_) is formed, and the monomer gets incorporated as part of the filament. (e) G-actin molecules in filaments are also attached to the grid with energy *E*_1_, mimicking the energy of the link between F-actin and anchor proteins in the inner plasma membrane, such as integrins (see Methods).

##### F-Actin depolymerization

(a) If the position [*x, y*] sampled is occupied, the probability of removing the G-actin is computed as *P*^−^ = *e*^−*E*^, with *E* being the sum of the energy of all links of this molecule; (c) At the back of the polar F-actin filament, depolymerization is favored by releasing the bond between the last G-Actin and the rest of the polymer (mimicking the destabilization of the G-Actin bonds at the pointed end), so the probability of removing the last G-Actin is only *P*^−^ = *e*^−*E*_1_^.

Results from numerical simulations of the model (plotted in Fig. 1 for three different time points, time lapse movie presented as Supp. Movie 1) show that F-actin formation occurs in three distinct phases. The first column corresponds to a **nucleation phase**, characterized by fast assembly and disassembly of short lived F-actin composed of very few monomers. In this phase, most of G-Actin is still at the cytoplasm (Fig. 1A), while the number of F-Actin (Fig. 1B) and the cortex occupancy (Fig. 1C) show an initial lag phase followed by cooperative polymerization.

**Figure 1:**
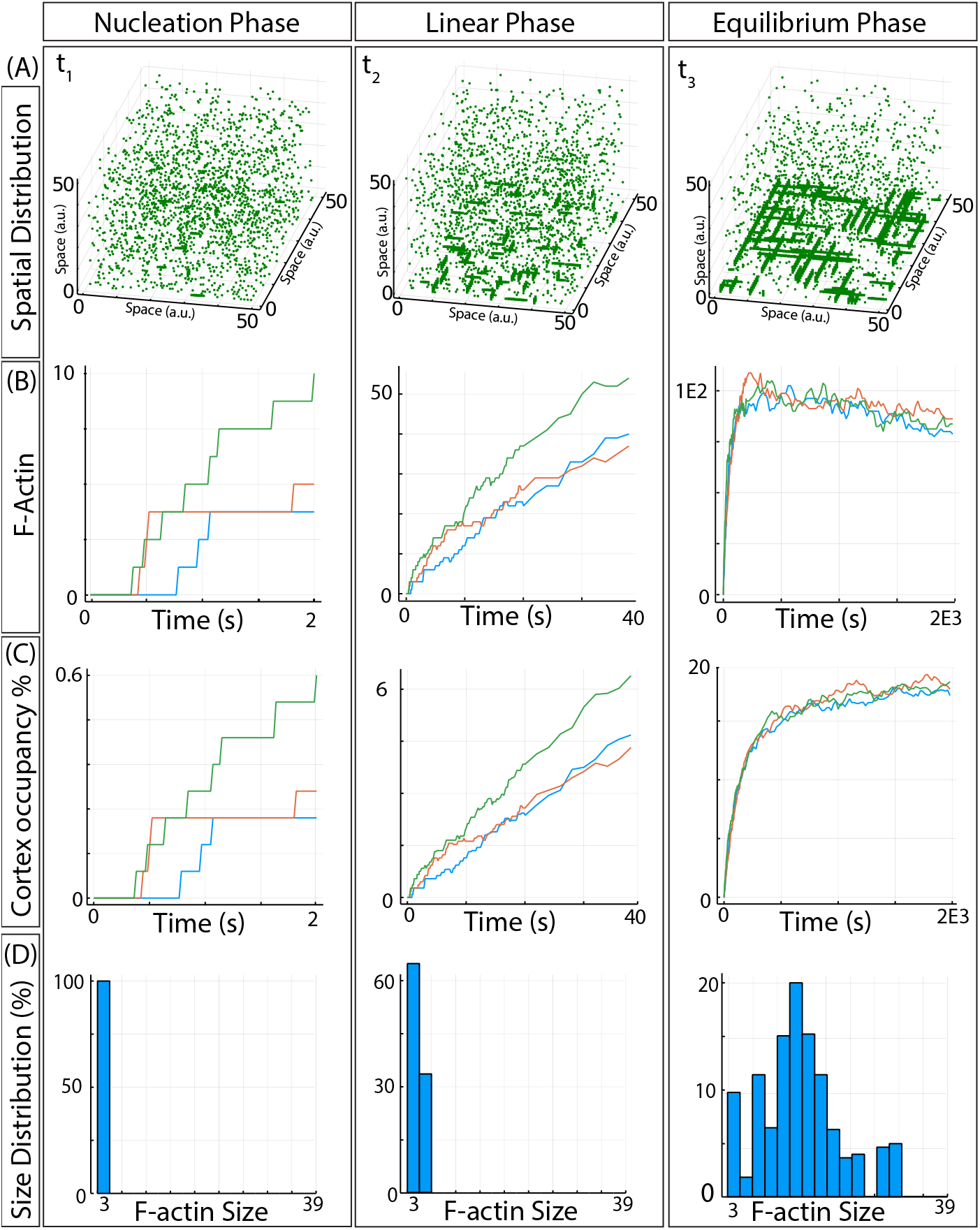
Dynamics of F-actin polymerization. (A) Snapshots of the system at three different simulation times, corresponding to the three regimes: nucleation, polymerization and equilibrium/reorganization. (B) Number of filaments forming as a function of time during the three regimes. (C) Percentage of the nodes in the network occupied by actin molecules as as a function of time during the three phases. (D) Probability size distribution of F-Actin for the three different phases.

The size distribution of F-actin polymers has been shown to strongly affect the fluidity and viscosity of polymer solutions, as well as in the cytoplasm, which in turn influences multiple important cellular properties and functions (Janmey *et al*, 1986; Trepat *et al*, 2007; Wang *et al*, 2019). The size distribution of the filaments (Fig. 1D) in this phase follows an exponential distribution, with many monomers being part of small filaments. This distribution, and the dynamics (lag phase and cooperative assembly) reproduce the experimental observations *in vitro* (Oosawa and Kasai, 1962; Oosawa and Asakura, 1975; Tobacman and Korn, 1983).

The second column corresponds to the **polymerization phase**, characterized by F-actin elongation (Fig. 1A), where the amount of F-Actin (Fig. 1B) and total G-Actin in filaments (Fig. 1C) increases at an almost linear pace. The size distribution is still monotonically decreasing, with F-Actin longer than in the previous phase, again in good agreement with experimental observations (Oosawa and Kasai, 1962; Oosawa and Asakura, 1975; Tobacman and Korn, 1983).

The final **equilibrium/redistribution phase** is characterized by the presence of large filaments and a few short-lived smaller filaments (Fig. 1A). The number of total molecules in filaments fluctuates around a constant value (Fig. 1C), while the number of filaments is slowly decreasing due to the gradual disassemble of these shorter polymers, to the expenses of the further elongation of the longer more stable filaments (Fig. 1B). The size distribution now shows a Poisson-type distribution, again in agreement with experimental observations (Oosawa and Kasai, 1962; Oosawa and Asakura, 1975; Tobacman and Korn, 1983).

Interestingly, and despite the simplicity of our model, these three phases (nucleation, polymerization and equilibrium/redistribution) reproduce the well-known dynamics and the size distribution of the three regimes observed experimentally (Fig 2 in Ref. Kuhlman (2005)).

**Figure 2:**
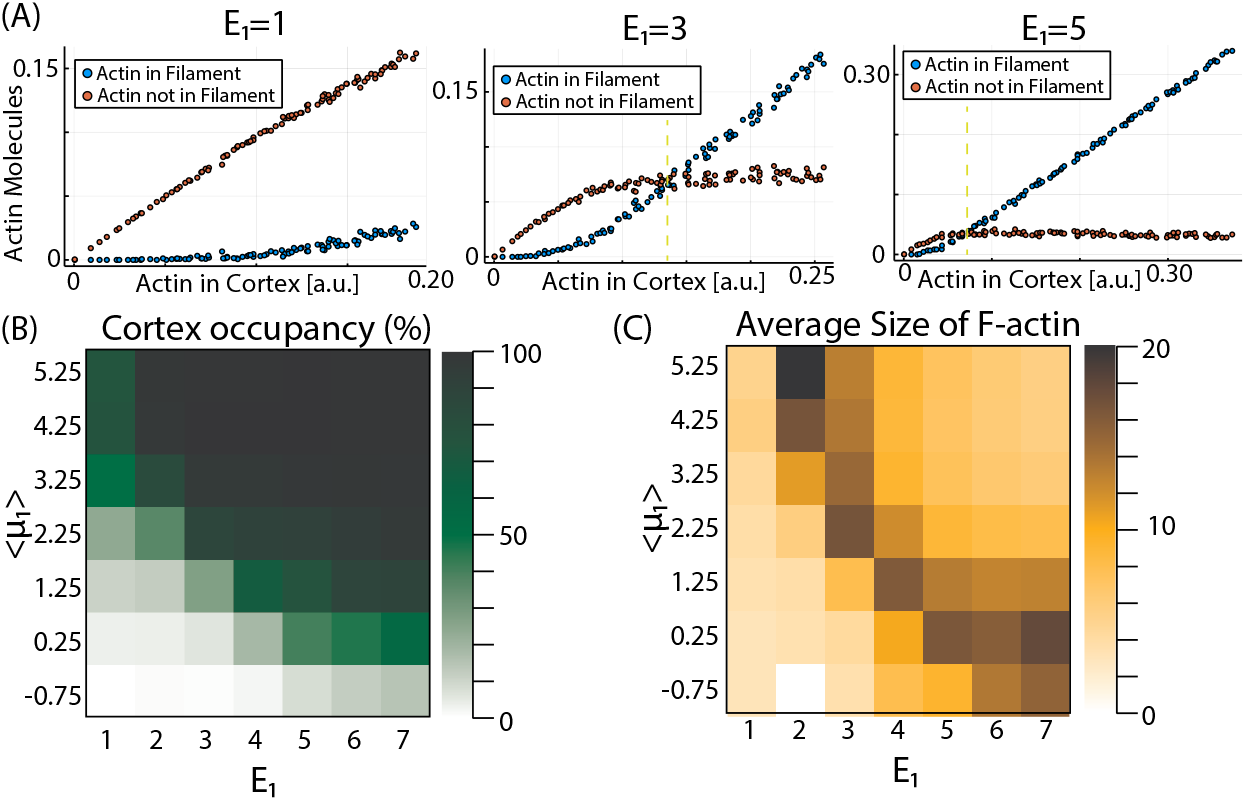
Dependence of F-actin features on model parameters. (A) Dependence of the amount of G-Actin in monomer and polymer configuration on the total amount of G-Actin in the system *N*_1,0_ for three different values of *E*_1_. (B) Phase diagram plotting the cortex occupancy for different values of the average <*μ*_1_ > and *E*_1_. (C) Phase diagram plotting the average length of F-Actin for different values of the average <*μ*_1_ > and *E*_1_.

### The model predicts a critical concentration of G-actin for polymerization

Another important observation from experimental work *in vitro* is shown in the early studies of Oosawa and Kasai (Oosawa and Kasai, 1962), where F-actin polymerization occurs only above a critical concentration in G-Actin in the system, to then increase linearly with the amount of available G-actin. After this transition point, the amount of monomers should remain constant and independent on the total levels of G-Actin in the system (all new G-Actin entering the system does so as part of F-Actin polymers). To test this in our model, we modulated the total number of G-Actin in the system (*N*_1,0_), and monitored the number and features of the F-actin formed at the cortex. Results are shown in Fig. 2A. The model predicts a critical value (yellow dashed vertical line) above which, the amount of total G-Actin in the cortex (blue dots) increases linearly, while the amount of G-Actin not in filament configuration (red dots) remains constant. This critical concentration is more evident for higher values of the energy *E*_1_ (right panel in Fig. 2A, lines drawn as a guide to the eye to illustrate the expected results in experimental observations).

### The model allows us to study the dynamics of F-actin for different experimental conditions

It is well known that some key properties of the inner membrane cortex depend on parameters such as concentration of G-actin and affinity towards the membrane, but these parameters are difficult to modulate *in vivo.* In this section, we take advantage of our framework to study these dependencies directly by changing parameters of the model and monitoring how the polymerization and depolymerization of F-actin is being affected. Results are shown in Fig. 2B-C.

At equilibrium, the model shows robust polymerization for a wide range of parameter values, with the total occupancy of the cortex (Fig. 2B) increasing as we increase the average value of the potential <*μ*_1_ > (that regulates polymerization) and the energy of the link *E*_1_ (that regulates depolymerization).

A more interesting result appears when the average size of F-actin is computed (Fig. 2C). We see that, for intermediate values of *E*_1_ and <*μ*_1_ >, the formation of F-actin is maximized, and large polymers dominate versus smaller polymers and monomers *N*_1,0_, in agreement with previous studies (Oosawa and Asakura, 1975). When we compoute the amount of G-Actin molecules distributed in F-actin of different sizes, we observe an exponential distribution (Supp. Fig. 1), as predicted in earlier studies (Edelstein-Keshet and Ermentrout, 1998).

In conclusion, despite the simplicity of our coarse-grained approach, the model is able to capture many key aspects of F-actin polymerization: its dynamics, the existence of a critical concentration of G-Actin, the size distribution in steady state and how it depends on monomer concentration. The phase diagrams also show that increasing <*μ*_1_ > is equivalent to increase <*E*_1_ > (phase diagrams are symmetric respect to the diagonal). This is true when monitoring the concentration of F-Actin (Fig. 2B) and its average size (Fig. 2C), but not for other features of F-Actin, as we will see later. Also, we can also observe that values around *E*_1_ = 5, <*μ*_1_ >= −1.25 result in the most efficient polymerization (i.e., many large filaments at low cortex occupancy).

### The model reproduces filament treadmilling

In conditions of constant concentration of G-actin, F-actin grows principally at the barbed end and shortens at the pointed end. When the rate of polymerization and depolymerization are balanced in a given F-actin, it produces an apparent displacement of F-actin known as treadmilling, which is essential for cell motility and other cellular processes (Lee and Dominguez, 2010; Pollard and Borisy, 2003).

To study the process of treadmilling in our model, we perform time lapse simulations where filaments can be tracked to monitor their position. Snapshots of the cortex at three time points of a given simulation are shown Fig. 3A, where we highlighted two filaments that move up to down (red) and right to left (blue). Filament treadmilling can be seen also during Supp. Movie 1.

**Figure 3:**
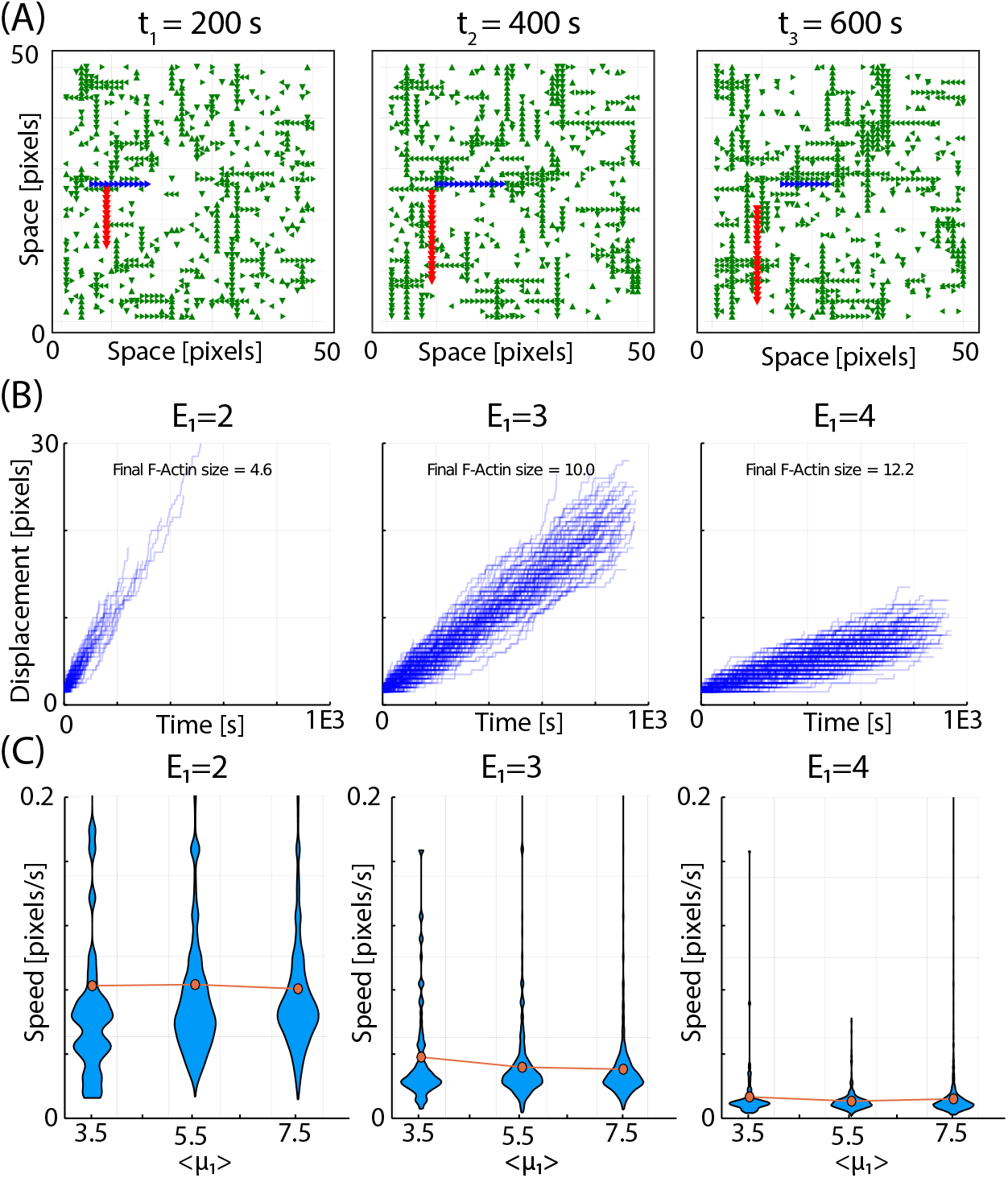
The model reproduces F-actin treadmilling. (A) Snapshots of the cell cortex showing two filaments (highlighted in red and blue) moving in different directions due to directional treadmilling. (B) Displacement of all filaments in a simulation showing the dispersion in velocity values for different values of energy *E*_1_. (C) Dependence on the velocity of filaments on the potential <*μ*_1_> for different values of the *E*_1_.

Treadmilling in our model occurs because polymerization is probabilistic, while depolymerization is time-dependent (i.e, the effective probability of disassembly increases with time). This way, at some average F-actin length, the rate of polymerization balances the rate of depolymerization, resulting in the apparent directional displacement of the filaments.

Fig. 3B plots the displacement overtime of the filaments for three simulations with different values of *E*_1_, showing that the average speed of the filaments depends strongly on this parameter.

Fig. 3C plots the speed of the F-actin for the three energies and for different values of <*μ*_1_>. Interestingly, the speed of treadmilling does not depend strongly on <*μ*_1_> (which strongly affects the size of F-Actin, as shown in the previous section). This means that changes in *E*_1_ modulate both F-Actin speed and length, while changes in <*μ*_1_> only affect the average length while maintaining the average displacement velocity.

Figs. 3B-C illustrates the strong variability for the speed of the different F-Actin in the same simulation. An analysis of the speed of the filaments and their length (Supp. Fig. 2), shows that the shortest filaments move faster that the average, while the speed of larger filaments does not depend on their length.

### The model reproduces cooperative network assembly

#### Box 2

##### Actin Crosslinkers (ACs) and network formation.

(a) A position in the grid [*x, y*] is sampled at random. (b) If the position is located between two parallel, anti-parallel or perpendicular F-actin polymers, the probability of insertion an AC molecule is *P*^+^ = *e*^*μ*_2_^. (b) When an AC is inserted, a link of energy *E*_2_ is established between the molecule and each of the G-Actin molecules; (c) When the position sampled is already occupied by an AC molecule, its removal is accepted or rejected using the Metropolis method (*P*^−^ = *e*^−2·*E*_2_^). Therefore, the strength of *E*_2_ can be modulated to produce stronger or weaker networks of Factin. (d) Removal of G-Actin that are linked to an AC molecule is now *P*^−^ = *e*^−*E*_0_−*E*_1_−*E*_2_^, so ACs also contribute to stabilization of the filaments, as it occurs *in vivo*.

In this section, we modify our framework to study the effect of Action Crosslinkers (ACs) in the configuration of the inner plasma membrane. Details of how ACs are modeled are explained in the Methods section. Snapshots of the system in the presence of ACs are shown in Fig. 4A for three different time points (time-lapse movie shown as Supp. Movie 2), showing an initial state with several small networks that eventually grow in size by fusion to other networks and by increasing the length of the F-actin.

**Figure 4:**
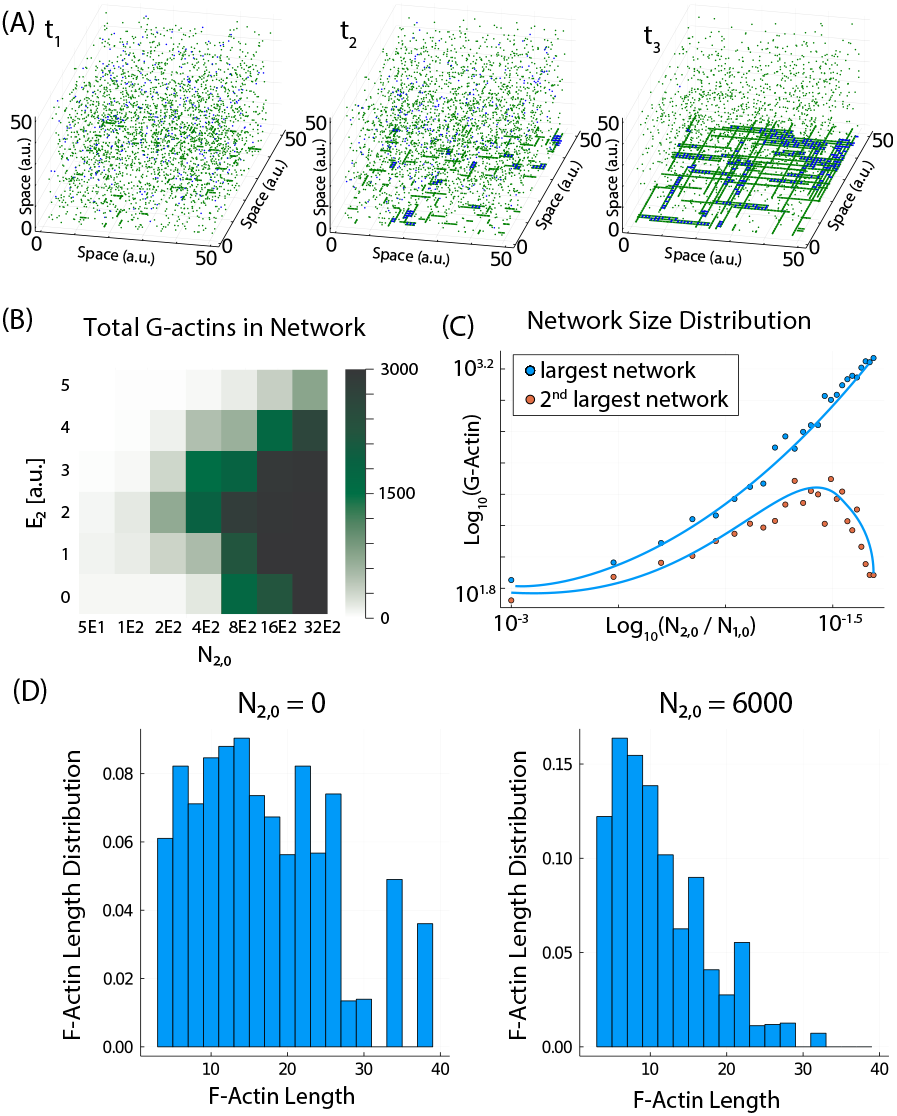
The model reproduces the dynamics of network formation. (A) Snapshots of system for different time points of a given simulation. ACs are added at *t*_1_. (B) Phase diagram showing the average network size at equilibrium for different values of *E*_2_ and *N*_2,0_. (C) Size of largest and second largest networks for different ratios of *N*_2,0_/*N*_1,0_. (D) Length Distribution of filaments without (left) and with (right) ACs.

It has been shown experimentally that the formation of F-actin networks and bundles (Bartles, 2000) only takes place above a critical concentration of ACs (Kierfeld *et al*, 2005, 2007; Ayscough, 1998). To study this, we computed a phase diagram (Fig. 4B) where the average amount of monomers in the cortex is computed for different values of the energy *E*_2_ and the total amount of of ACs (*N*_2,0_). The plot shows that, for a given value of the energy *E*_2_, the formation of networks only occurs above a critical value of the concentration of ACs, in agreement with experimental observations (Kierfeld *et al*, 2005, 2007; Ayscough, 1998). Interestingly, for a given concentration of ACs, the size of the network is maximized at intermediate values of the energy *E*_2_.

Previous studies also focus on the competition between networks for different amounts of ACs (Alvarado *et al*, 2013), showing that the largest network increases at the expenses of the decrease of the second largest network when the amount of ACs is increased. This competition also takes place in our model (Fig. 4C), in agreement with the previous studies (Alvarado *et al*, 2013).

Network formation is a highly cooperative mechanism (Shin *et al*, 2009) (as the network grows, the probability of successful adding more ACs also increases). The dynamics of incorporating the molecules into the networks (Supp. Fig 3) shows an initial very fast incorporation of ACs and G-actin as soon as the ACs are added into the system.

Another important experimental observation measures the reduction in the width of the length distribution of F-actin after addition of ACs (Biron and Moses, 2004). Our model also shows the same reduction of the standard deviation in the distribution, in the presence of ACs (Fig. 4D).

In conclusion, our model reproduces many key experimental features of F-actin networks, such as the cooperative assembly, the threshold concentration of ACs, the competition between networks and the homogenization of the length of the filaments in the presence of ACs.

### The model reproduces actomyosin oscillations

In this section, we use our GC framework to study the dynamics of actomyosin when we include tension induced-disassembly of the F-actin, mediated by the mechanical action of Myosin. Details of how Myosin and F-actin load is introduced in the model are explained in the Methods section and summarized in Box 3. Snapshots of the system at three time point of the same simulation are shown in Fig. 5A (time-lapse movie shown in Supp. Movie 3). In the first time point (left panel), a large percentage of the molecules are at the cortex, as part of a network. Then (central panel), the network is disassembled and the molecules are mainly at the cytoplasm. The next time point (right panel), shows the actomyosin cortex starting to form again.

#### Box 3

##### Myosin incorporation and release

(a) A position [*x,y*] in the grid is sampled, (b) If the position is empty and located between two anti-parallel F-actin polymers, the probability of insertion of a Myosin is *P*^+^ = *e*^*μ*_3_^. (c) When an Myosin is inserted, a link of energy (*E*_3_) is established between the molecule and each of the G-Actin molecules. (d) Removal of G-Actin linked to a Myosin molecule is now *P*^−^ = *e*^−*E*_0_−*E*_1_−*E*_3_^, so Myosin molecules also contribute to stabilization of the filaments.(e) When the position [*x,y*] sampled is already occupied by a Myosin molecule, its removal is accepted or rejected using the Metropolis method (*P*^−^ = *e*^−2·*E*_3_^).

##### Myosin induced network disassembly

(a) Myosin molecules are assumed to produce a power *W*_3_, measured in units of [kT/s], which produces a load *F* in the F-actin attached. (b) When the accumulated load in a given F-Actin exceeds the energy of all the bonds that maintain it attached to the inner plasma membrane (*E_t_* = *E*_1_ × *L*, being *L* the length of this F-actin), the filaments can detach. (c) Detached F-Actin depolymerizes and monomers are back to the cytoplasm. (d) Since all F-actin in a network are attached via ACs, once a F-actin a small percentage of the load *δF* is transmitted to the rest of the F-Actin (see Methods for a more detailed explanation).

**Figure 5:**
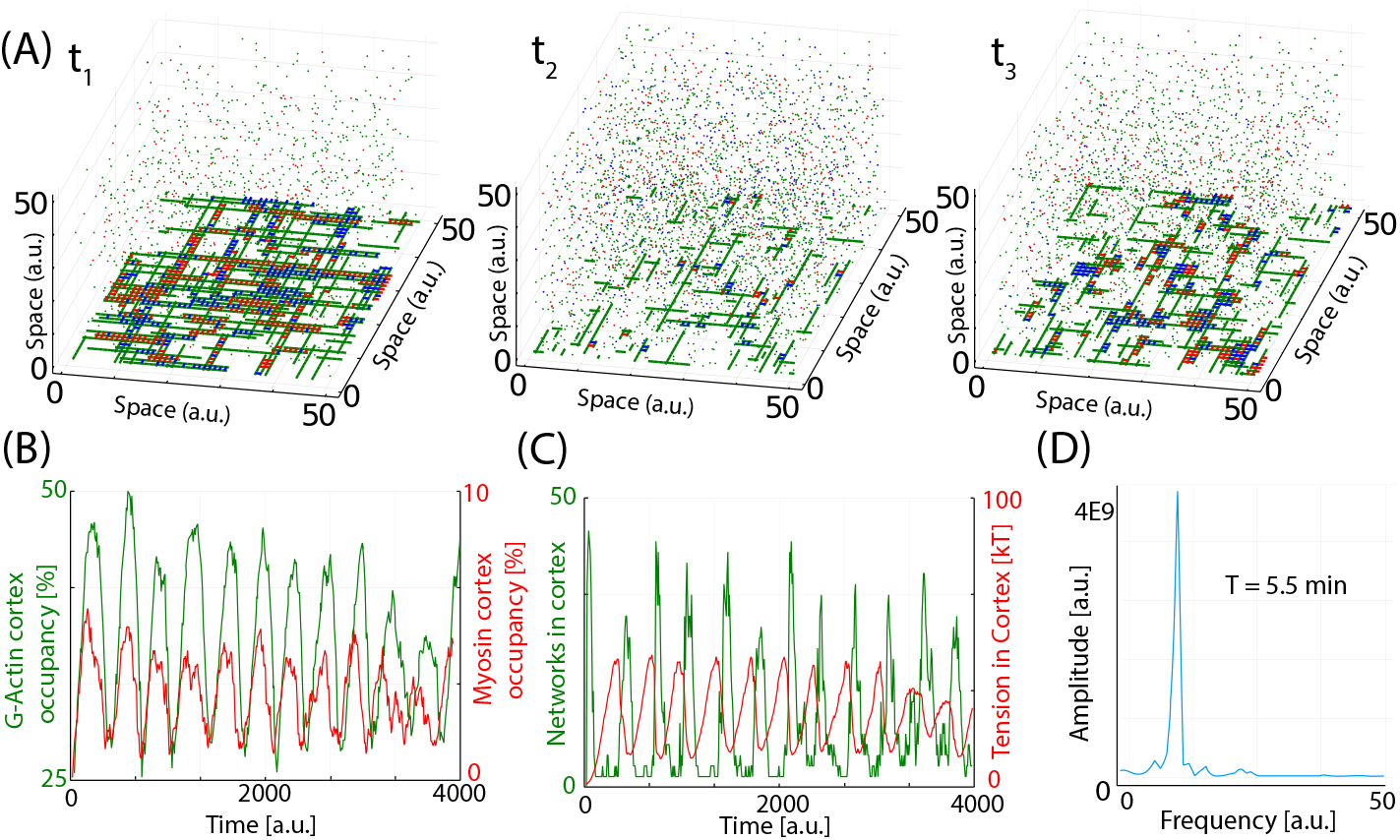
The model reproduces actomyosin oscillations. (A) Snapshots of the system for three different time points of a given simulation (green is G-Actin, blue is ACs, red is Myosin). (B) Temporal evolution of total actin and myosin in the cortex. (C) Temporal evolution of tension (red) and number of bundles (green).

The total amount of G-Actin (green) and Myosin (red) in the cortex during these oscillations are plotted in Fig. 5B, where we can see periodic changes in both levels in the form of autonomous oscillations with a defined period. Fig. 5C plots the total tension in the cortex (red line) and the number of simultaneous networks (green line). Interestingly the tension increases in anti-phase with number of networks (i.e., as the largest network takes over the system, the tension increases until some of the filaments reached their threshold in tension). The release of tension is also much faster that the disassembly of the network, and the difference between these two time scales is at the core of this oscillatory dynamics. Fig. 5D plots the Fourier transform of the temporal evolution in the number of G-Actin at the cortex, with a clear peak evidencing the periodic behavior. The period of oscillation is around 5.5 minutes, with is similar to the values measured *in vivo* (Solon *et al*, 2009; Sedzinski *et al*, 2011; Machado *et al*, 2014; Dierkes *et al*, 2014; Koride *et al*, 2014).

Previous experimental studies (Valencia-Expósito *et al*, 2016) shown that actomyosin oscillations accelerate, decrease in amplitude and become less periodic and more stochastic when the amount of active Myosin is increased. To test this situation, we compared in Supp Fig. 4 the oscillations for low (*M*_3,0_ = 200, panel A) and high (*M*_3,0_ = 4000, panel B) Myosin. We can see that, as we increase Myosin, oscillations are smaller, faster and more stochastic, in agreement with the experimental observations (Valencia-Expósito *et al*, 2016).

### The attachment of Actin molecules to the cortex modulates the network architecture and dynamics

In previous sections, we have corroborated that the model reproduces many already established important aspect of the actomyosin machinery. In the present section, we proceed to test the model and its ability to predict biological responses to changes in key parameters of the system, such as the anchor of F-actin to the cortex. By changing the values of the attachment strength of the Actin molecules to the cortex *E*_1_, our model predicts a high dependence of the F-Actin architecture and network dynamics on the attachment strength of the Actin molecules to the cortex (Fig. 6A-B). Our results show that increasing *E*_1_ results in a more dense cortex with F-Actin of larger length (Fig. 6A, snapshots taken at the maximum of the oscillation). Focusing on the oscillatory dynamics, both amplitude and period of the oscillations increase when *E*_1_ is increased (Fig. 6B).

**Figure 6:**
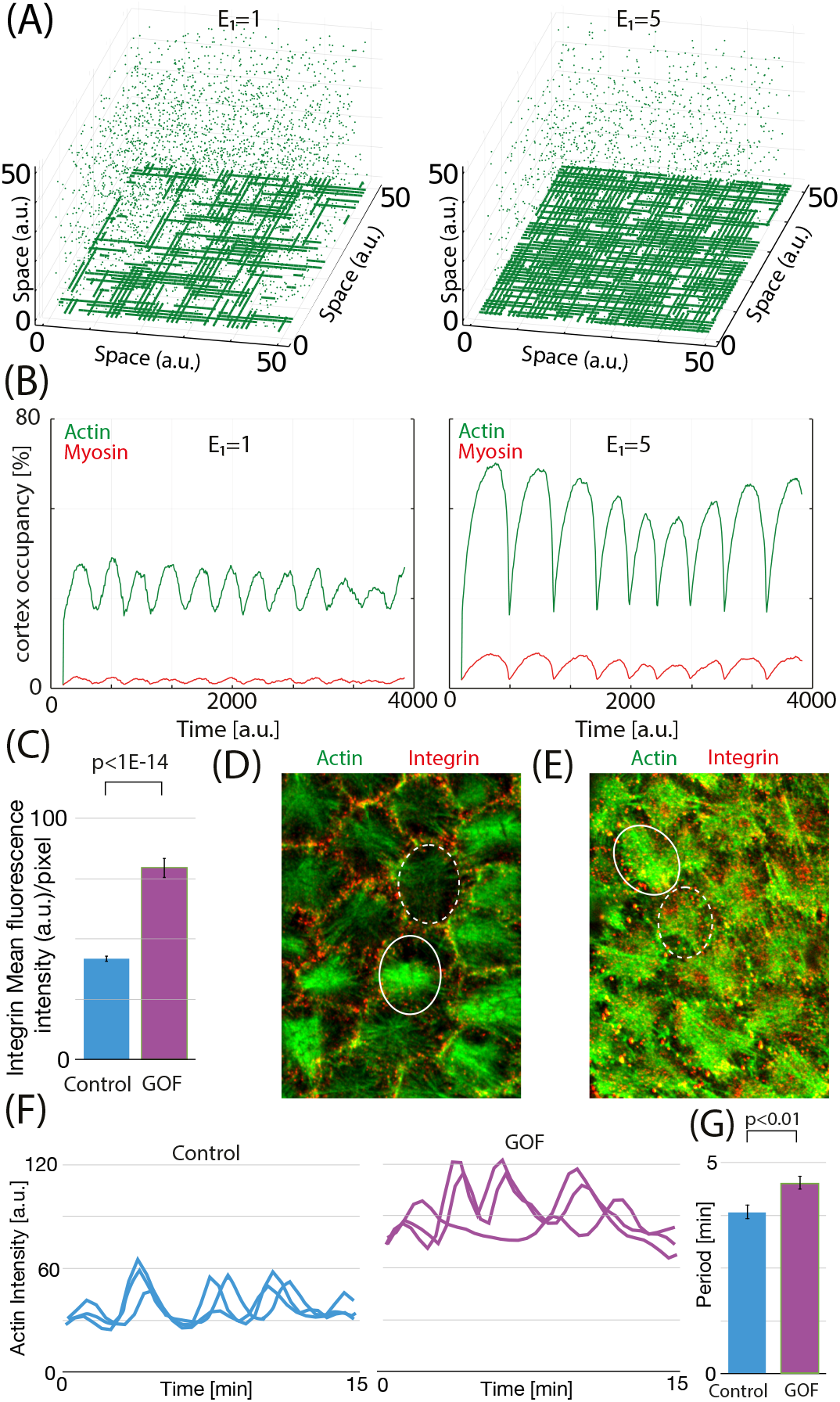
The model predicts the effect of integrin overexpression. (A) Snapshots of the system for two different values of *E*_1_ (green is G-Actin, snapshot taken at the point of maximal value of occupancy in a oscillation). (B) Temporal evolution of total actin and myosin in the cortex for the two values of *E*_1_ tested. (C) Mean fluorescent intensity of Integrins in stress fibers of follicle cells in control and *α*PS1; *β*PS overexpression (labeled as Gain Of Function (GOF)) conditions. (D-E) Confocal images of the basal surface of follicle cells for control (D) and GOF (E) conditions expressing the F-actin live marker Ubi-LifeActinGFP stained for anti-GFP (green) and anti-BPS integrin (red). Circles denote follicle cells with the highest (solid line) and lowest (dashed line) levels of F-actin fluorescence respectively. (F) Quantification of the dynamic changes of basal Factin in three individual follicle cells for control (left) and GOF conditions. (G) Basal F-actin average oscillation period for control and GOF conditions.

Integrins are proteins known to mediate attachment of F-actin to the cortex through actin binding molecules, such as Talin (Calderwood *et al*, 2000). Using the actomyosin fibers present on the basal side of *Drosophila* follicle cells as a well-established model to study actomyosin dynamics (He *et al*, 2010), we and others have previously shown that integrins are required for the correct recruitment, formation and maintenance of these basal F-actin fibers (Qin *et al*, 2017; Santa-Cruz Mateos *et al*, 2020).

To test whether the predictions made from the model hold when applied to *in vivo* systems, we analysed the consequences of increasing the amount of F-actin fibers at the cortex. To do this, we decided to increase integrin levels in follicle cells (FCs).

Integrins are trans-membrane proteins that mediate cell-to-matrix and cell-to-cell adhesion, with important roles in cell adhesion, migration, proliferation and survival (Brakebusch and Fässler, 2003). Integrins are know to be coupled to Actin molecules in the cortex through physical linkage (Calderwood *et al*, 2000), and they have been shown to shape the architecture and mechanical properties of actomyosin networks (Santa-Cruz Mateos *et al*, 2020). Integrins are heterodimeric proteins composed of an *α* and a *β* subunit. Two integrins are expressed in follicle cells of *Drosophila*: the *α*PS1*β*PS (PS1) and the *α*PS2*β*PS (PS2). Since the transport of integrins to the surface requires both subunits of the integrin protein, to increase integrin levels in follicle cells, we overexpressed both *α*PS1 and *β*PS subunits simultaneously using the UAS/Gal4 system (Brand and Perrimon, 1993), so we can compare the dynamics and configuration of the actomyosin cortex in conditions of control and *α*PS1; *β*PS overexpression (labeled as Gain Of Function, GOF).

We found that over-expression of the PS1 hetero-dimer in all follicle cells, using the traffic jam Gal4 line (tj-Gal4, (Li *et al*, 2003)) driver (tj>PS1), resulted in an increase in the amount of integrin present in the actomyosin filaments (Fig. 6C).

Moreover, we found that when the PS1 integrin was overexpressed in all FCs, the amount of F-actin accumulated in fibers was higher and more stable than that found in control follicle cells (Fig.6E and G). These results suggest that increasing the amount of integrins in FCs is an efficient way to increase the amount of F-actin fibers at the cortex, similar to what happens in the simulations when increasing the value of *E*_1_. Interestingly, we found that F-actin filaments of FCs overexpressing PS1 appear longer than in wild type conditions, in full agreement with the model predictions (Fig. 6A, E).

To test the effects of elevated integrin levels in the oscillatory behavior of F-actin, we performed live imaging of egg chambers for both control and GOF conditions expressing the *in vivo* marker for F-actin, Ubi-LifeAct-GFP (see Material and Methods and ref (Santa-Cruz Mateos *et al*, 2020)). We found that the basal F-actin of tj>PS1 FCs underwent periodic fluctuations (n=8, Fig. 6G, Sup. Movies 4 and 5). However, in contrast to the marked and periodic changes observed in control FCs, the oscillation period in tj>PS1 FCs showed a high degree of variability, and both maximum and minimum (solid and dashed circles, respectively) show higher intensity levels in GOF conditions, compared to control (Fig. 6D-E). This suggests that, in GOF conditions, the actomyosin complex is not being fully disassembled during the course of the oscillation, compared to a more complete disassemble in control FCs. Quantification of the pulsation frequency revealed that the period of tj>PS1 FCs was 17 % higher than that of controls, in agreement with model predictions.

In conclusion, our model does not only reproduces well known features of the actomyosin cortex, but can also make robust predictions, as it reproduces the features of basal F-actin oscillations in wild type and in experimental perturbations.

## Discussion

In this paper, we presented the the first GC model of the actomyosin cortex formation and dynamics, that includes interactions between molecules and with the inner plasma membrane. The most complex task in the design of a useful GC model is to select the main relevant ingredients that govern the dynamics of the system. The full version of the model is based on only three basic aspect of the actomyosin machinery: (1) polymerization and depolymerization of F-Actin from G-Actin at the inner cellular membrane, (2) supramolecular organization of F-Actin in networks mediated by ACs, (3) reorganization of the network mediated by myosin. Despite this highly simplistic approach, our model successfully reproduces many of the most relevant processes involving F-actin: (1) FActin formation occurs in three regimes (nucleation, linear and equilibrium/redistribution); (2) size distribution in these three regimes; (3) threshold for F-actin polymerization (4) treadmilling; (5) competition between networks; (6) cooperativity and periodic assembly and disassembly of actomyosin networks.

The model assumes several important simplifications. It assumes that the inner plasma membrane is a two dimensional grid where F-actin molecules of the same orientation are not allowed to overlap (cortex thickness has been estimated to be around a few hundred nm (Chugh *et al*, 2017), while diameter of F-actin is around 10 nm). In addition, F-actin is often composed by many more G-actin subunits than in our model. Also, we have not incorporated the fact that F-actin can grow from both barbed and pointed ends, but the dynamics of growth in the barbed end is much faster than in the barbed end. This can be easily implemented as an extension of our model, although we expect that the conclusions derived from the model will be equivalent.

The model also assumes that molecules in the reservoir diffuse instantaneously compared to two dimensional diffusion while in the cortex. In addition, G-Actin does not diffuse while in the cortex, and it is assume to be linked to the membrane. Another simplification is to assume ATP in excess in the reservoir, so the effect of ATP is not explicitly included in the model for simplicity. In the same direction, we do not model explicitly the interaction between Myosin and ATP. Also, the grid only has two directions, which is a strong simplification compared to the experimental situation. A potential extension of the model is to allow another direction by setting up the grid as an hexagonal lattice.

Our model shows that treadmiling depends strongly on the energy of the link *E*_1_ (which mainly controls depolymerization) and not with the value of *μ*_1,0_ (which mainly controls polymerization). Also treadmiling in our models is nondirectional, contrarily to the directional dynamics observed in filopodia and lamelopodia (Pollard and Borisy, 2003). This is due to the lack of regulatory proteins in our model, such as ADP/cofilin and caping subunits that break the symmetry and favor directional growth of the filaments.

Several studies show that the architecture of the network ultimately determines the dynamics of the interaction of actin with myosin (Reymann *et al*, 2012). This architecture depends on the different types of ACs that are known to link F-actin. Implementing other crosslinkers with other binding characteristics will result in different scenarios, and will be part of a follow up study to this one.

Finally, the dynamics of polymerization of nonmuscle Myosin is also a complex and multi-step process that it is not fully characterized. For simplicity, our model simply assumes a constant concentration of Myosin already in its filament form.

All these limitations can be easily addressed in a more detailed version of the model, using high performance computers, but we believe that the relevance, impact and usability of a model is higher when the main features can be obtained with conventional hardware. In addition, the model is not designed to reproduce the small details of the actomyosin cortex, but to illustrate the sets of minimal interactions that set the most important characteristics at the systems level.

When comparing to the experimental data in the last section of the results, the experiments show increased stochasticity in the oscillatory dynamics when integrin levels are increased. This is not the case in the model, where a single network is formed in every oscillation, and oscillations are very periodic independently on the level of *E*_1_, at least for these parameter values. This highlight one of the clear limitations of the model, since in the real systems many actomyosin subnetworks are formed simultaneously in these conditions, resulting in a less regular behavior (the same occurred when modulating the myosin levels experimentally (Valencia-Expósito *et al*, 2016)). Also, the experiments shown F-actin of length larger than the average cell diameter, so maybe the stochasticity arises from coupling between cells. These qualitative observations, and the potential mechanical coupling between cells remain to be studied in detail in the future.

## Conclusion

The study of the fundamental properties of transiently cross-linked networks is highly complicated due to the multiple space and time-scales involved (Huber *et al*, 2013). Systems such as the actomyosin cortex, where features emerge due to interactions between many players as systems properties, present an optimal problem that can be approached using GC models. One limitation of these CG models is that, due to the minimal set of players and interactions, they are not suited to study details of the biological system in question. On the other hand, they are very useful to isolate the most important players and interactions that are at the core of basic biological phenomena (Ruiz-Herrero *et al*, 2013).

In our model, Myosin motors and cross-linkers interplay with F-Actin polymerization and affect the dynamics of the network, its conformation, its dynamics and its mechanical properties. Our model can be used to study the dynamics of polymerization, the basis of treadmiling, the cooperative phenomena that drive polymerization and network formation, the thresholds and critical concentrations. In addition, our model allows us to study how adhesion is shaping actomyosin networks in the cell cortex, resulting in periodic oscillations that reproduce many aspects that have been observed experimentally. model can allow us to study how tension is shaping actomyosin networks in the cell cortex, resulting in periodic oscillations that reproduce many aspects that have been observed experimentally. Our framework shows that, despite the complexity of the actomyosin cortex, many of its key features can be explored and reproduced from very few well-characterized interactions at the molecular level.

## Materials and Methods

### Modeling framework and model assumptions

Numerical simulations of the actomyosin cortex are performed using a finite multi-compartment GC framework developed in house (code available as supplementary material). In this system, the inner cellular membrane is defined as a two dimensional region where total energy and number of molecules are not fixed, so it can be considered an open system and, therefore, the probability distribution of the different micro-states of the systems is represented statistically by a Grand Canonical ensemble. Moreover, since several types of particles coexist and interact (in our case: G-Actin, Actin Crosslinkers (ACs), and Myosin), a semi-grand canonical approach is used (Madurga *et al*, 2011; Ueno and Shibuta, 2018).

The code is written in Julia language (arXiv:1209.5145), and the following libraries have been used: LightGraphs, FFTW, Plots, Random, Printf, Dates, Statistics, StatsPlots, KernelDensity, GLM.

The model is solved numerically using a Monte-Carlo (MC) computational algorithm that relies on repeated random sampling to obtain numerical results. The dynamics is determined by the Metropolis method (Metropolis and Ulam, 1949), used to evaluate if a given transition between an “old” and a “new” state of the ensemble is accepted. The algorithm proceeds as follows. First, a position [x,y] at the cortex grid is chosen at random. Next, a change in state in the system due to an insertion or removal of a molecule in this particular position is evaluated. This process is then repeated for each of the three particles and at every time step. In the following paragraphs we explain the biological motivation behind the design of the framework, and how this features are implemented at the computational level.

#### Cortex

The actomyosin cortex is a pseudo-two dimensional mesh of actin filaments actin crosslinkers and myosin attached to the inner plasma membrane by anchor proteins. The contractile tension imposed by myosin motors in the network drives important changes in the shape of cells (Chugh *et al*, 2017). Our model represents this inner plasma membrane as a two-dimensional grid of fixed size where molecules can interact (see Supp Fig 5A for an illustration of the system). Each node of the grid can accommodate one G-Actin molecule in vertical orientation (north, south) and one in horizontal orientation (east or west), to allow overlapping of perpendicular filaments. Bonds between molecules are simplified as contact interactions (i.e, molecules can interact if they are in neighboring positions of the grid). For simplicity, molecules are not allowed to move or rotate when in the grid. The strength of the interactions is modulated to produce different scenarios. G-Actin molecules occupy the center of pixels of the grid, ACs and Myosin are occupy the space in between pixels, linking adjacent monomers in neighboring pixels.

#### Cytoplasm

Directly attached the two dimensional grid, the model assumes a finite threedimensional reservoir, which mimics the role of the cytoplasm, where molecules diffuse freely (Supp Fig 5A). For simplicity, diffusion or movement in the cytoplasm is assumed to be much faster than the process of attaching and detaching to the cortex. Therefore, the cytoplasm is simply modeled as a zero-dimensional reservoir. The total number of molecules of each type in both regions (cortex+cytoplasm) is maintained constant throughout the simulation.

#### Transitions from Cytoplasm to Cortex

During the simulation, three types of molecules (i=1 for G-Actin, i=2 for ACs, i=3 for Myosin) move from the cortex to the cytoplasm (Supp Figs 5B-C) based on the value of a chemical potential *μ_i_,* computed as a generic function with the following form.

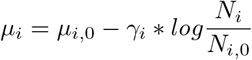

where *μ*_*i*,0_ is a reference potential for each molecule type, *N*_*i*,0_ is the total number of molecules of type *i* in the system, *N_i_* is the number of molecules of type *i* in the grid at a given time point. Parameter *γ_i_* modifies the shape of the function but does not affect the equilibrium of the system.

Therefore, values of *μ_i_* higher than zero favor the transition of a molecules of type *i* from the cytoplasm to the cortex, while values lower than zero disfavor it. Plots of the potential function for different values of *μ*_*i*,0_, *N*_*i*,0_ and *γ_i_* are illustrated in Supp Figs 5D-F.

When the position [*x,y*] being evaluated is empty, the probability of successful insertion of a molecule of type i into this position is calculated as *P*^+^ = *e*^*μ_i_*^, and the Metropolis method is used to accepted or rejected the change (a random number *P_rand_* from a uniform distribution between 0 and 1 is compared with the value of *P*^+^. The change in the state of the system is accepted if *P_rand_* < *P*^+^).

#### Transitions from Cortex to Cytoplasm

When the position [*x, y*] being evaluated is occupied by a molecule of type *i*, then the probability of removal from the cortex to the cytoplasm is calculated based on the energy *E* of all the links between this particular protein of type *i* and the other proteins of the cortex, calculated as *P*^−^ = *e^−E^*. Again, the Metropolis method is used to accepted or rejected the change.

#### Time calibration

To calibrate the temporal scale of the model, we use *in vivo* measurements of the polymerization of F-actin in cortex of mammalian cells (Fig. 1C in ref Murthy and Wadsworth (2005)), where authors estimated the time to reach equilibrium in the F-actin networks around 18 min. This value is used as a reference to establish a correlation between time iterations and real units of time, to be able to compare with experimental data.

In addition, the speed of the simulations in these type of systems depends by definition also on the size of the grid (due to the random sampling strategy). Therefore, the time is also calibrated by dividing it directly by the area of the two square grids (vertical and horizontal).

#### F-Actin polymerization and depolymerization

G-actin is a globular multi-functional and polar protein present in all eukaryotic cells. In the presence of ATP, G-actin can polymerize to form F-actin, a semi-flexible directional micro-filament composed of several thousand actin monomers (Carlier, 1991). F-Actin polymerization is mainly directional, i.e., new G-Actin monomers are incorporated at one end of the filament (barbed end), while G-Actin is released from the other end (pointed end), driven by the ADF/cofilin (AC) family of proteins. monomers (Edelstein-Keshet and Ermentrout, 1998).

Polymerization occurs when a G-Actin monomer enters the grid in front of another G-Actin or F-actin and with the same orientation (a link is established with energy *E*_0_). In addition, Actin molecules are maintained at the cortex by links to anchor proteins at the inner membrane layer, such as integrins (Santa-Cruz Mateos *et al*, 2020). Our model incorporates this feature by assuming that, apart from the links between monomers with energy *E*_0_, that G-Actin molecules in a filament are also linked to the inner plasma membrane with an energy *E*_1_.

Depolymerization at the pointed end occurs due to destabilization of the link between the last G-Actin of an F-actin and the rest of the filament. Therefore, the last G-actin is maintained at the cortex only by the link *E*_1_. For simplicity, G-Actin monomers that are not part of stabilized filaments are considered not to be linked to the inner membrane.

Experimental studies estimate the bonds mediated by ATP hydrolysis on the order of 30 kJ/mol (RT = 2.47 kJ/mol), we can estimate that *E*_0_ ≈ 12 in units of RT. This makes the probability of removing a G-Actin inside a F-actin (*P*^−^ = *e*^−*E*_0_−*E*_1_^) around 5 orders of magnitude less probable than removing the last G-actin in a filament (*P*^−^ = *e*^−*E*_1_^). Based on these estimations, the probability of breaking a filament as zero in this version of the model, for simplicity.

Polymerization and depolymerization of F-actin in our model is assumed to occur only at the cortex, and the steps are summarized in Box 1 in the main text. Figures related to F-Actin polymerization are plotted based on the effective value of the potential function, defined as:

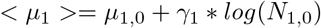

Previous studies define the nucleus for F-actin polymerization as the minimum number of aggregated monomers that is favored to grow (Oosawa and Kasai, 1962; Oosawa and Asakura, 1975; Tobacman and Korn, 1983). Based in this definition, the polymerizing nucleus in our model is composed of three monomers (the bond of the last G-Actin is destabilized), and therefore, F-Actin is defined when formed by three or more G-Actin monomers. This is in agreement with recent experimental observations Levin *et al* (2020).

#### Actin Crosslinkers (ACs)

F-actin molecules can cross-link or aggregate to form the actin cytoskeleton in the form of networks or bundles (Carlsson, 2010). This aggregation is mediated by Actin Crosslinkers (ACs) such as fascin, the Arp2/3 complex, Fimbrin, Filamin and *α*-actinin (Pantaloni *et al*, 2001; Mitchison and Cramer, 1996; Carlier *et al*, 2003; Sjöblom *et al*, 2008).

In brief, ACs establish and maintain bundles and networks of F-actin, introducing inter-filament attraction via a reversible interaction between one crosslinker and two actin monomers in a filament. The binding of ACs stabilizes the F-actin and drives the formation of structures of parallel and anti-parallel F-actin in the form of organized bundles or networks, which are at the core of many biological processes involving actomyosin, such as cell shape and cell movement. The concentration of ACs has been shown to dramatically alter the elastic properties of bundles and networks (Gardel *et al*, 2004). The process of insertion or removal of ACs in our model is explained in Box 2.

#### Myosin

Myosin motors are the major driving force in cell motility, muscle contraction and transport at the intra-cellular level (Pollard and Borisy, 2003). This highly conserved family of molecular motors are virtually present in all eukaryotic cells. Among them, myosin II (also known as conventional myosin) is the responsible for the generation of mechanical force in actin-based cell motility (Veigel and Schmidt, 2011).In nonmuscle cells, active myosin is commonly found as bipolar filaments (anti-parallel arrays of myosin molecules) of 10 to 30 myosin II dimers into filaments. These myofilaments interact with F-Actin by hydrolyzing ATPs and transforming the chemical energy into mechanical force, allowing their heads to tether F-actin and induce movement, contraction and tension (Mizuno *et al*, 2007) in the network (Cooper, 2000; Vogel *et al*, 2013) and in the cell membrane (Mason *et al*, 2013; Reversi *et al*, 2014).

The effect of Myosin depends strongly on the orientation of two F-actin molecules that it is attached (summarized in Supp Fig 6). In brief, if the F-Actin molecules are parallel (panel 1), the mechanical action of Myosin results simply in translocation of the molecular motor across the filaments. If the F-Actin molecules are anti-parallel (panel 2), the mechanical action of Myosin is known to induce sliding (Takiguchi *et al*, 1990) of the two F-actin molecules relative to each other.

F-actin molecules in the cortex are attached to the inner cell membrane, and therefore cannot slide freely (panel 3). In this conditions, the mechanical force is translated into tension in the filaments and in the network, affecting the shape of the membrane (Carvalho *et al*, 2013). Tension in the actomyosin has been shown to induce network reorganization (Houdusse and Sweeney, 2016), network disassembly (Wilson *et al*, 2010; Alvarado *et al*, 2013) and F-actin ruptures (Houdusse and Sweeney, 2016) has been studied extensively and have been shown to have a pivotal role in normal cell division (Sedzinski *et al*, 2011), in developmental disorders (Martin *et al*, 2010), and in cell motility (Wilson *et al*, 2010; Vogel *et al*, 2013; Haviv *et al*, 2008). Moreover, this sensitivity of F-actin to tension is at the core of a highly dynamic tension sensor in cells (Galkin *et al*, 2012; Hayakawa *et al*, 2011).

Based on these biological features, myosin and its effect in F-actin networks is introduced in the model as: (a) Myosin enters and exits the cortex, and binds to F-actin similarly to ACs (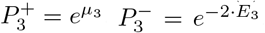, being *E*_3_ the energy of the bond between Myosin and one G-Actin molecule); (b) Myosin molecules are assumed to produce a power *W*_3_, measured in units of [kT/s], and induces a load *F* in the form of tension when linked to anti-parallel F-Actin molecules in a time dependent manner; (c) above a certain threshold, the tension induces direct detachment of the F-actin from the inner part of the membrane (Brugués *et al*, 2010); (d) A percentage of the tension supported by this F-Actin is redistributed to the other F-actin in the same network; (e) detached F-actin in the cytoplasm depolymerizes, increasing the amount of G-actin monomers that are available to enter the cortex (panel 4 in Supp Fig 6).

Since ATP is assumed in excess, and a single Myosin can hydrolyze many ATPs (and therefore, produce many mechanical effects on the molecules linked to it), our model simplifies the mechanical action of Myosin over F-actin as time dependent (i.e., tension accumulates while the Myosin as long as Myosin remains attached to the filaments).

In addition, the threshold tension that a given F-actin can support is assumed to depend on the strength of its attachment to the inner membrane and the extra-cellular matrix. This attachment is mediated by the integrin family of cell adhesion molecules (Brakebusch and Fässler, 2003), that are linked to F-actin by specific actin-binding proteins, such as talin, vinculin *α*-actinin and others (Calderwood *et al*, 2000). Therefore, since our model assumes that G-actin in the filaments are linked to the cortex with an energy *E*_1_, the threshold value before full detachment for a given F-Actin is computed as the sum of all its links (i.e., *E_threshold_* = *E*_1_ × *L*, being L the length of the filament).

Finally, the effect of redistribution of tension in the network is based on the fact that all F-Actin in a network are connected to the inner plasma membrane, which acts as an elastic load against network compression. When one of the F-Actin “cables” that support this load is removed from the network, a percentage of the elastic load is now being supported by the rest of the filaments of the network, that experiment an increase in the tension that they are supporting.

## Experiments

### *Drosophila* stocks and genetics

UAS-*α*PS1;UAS-*β*PS (Martin-Bermudo *et al*, 1997), the follicle stem cell driver traffic jam-Gal4 (tj-gal4, (Tanentzapf *et al*, 2007)) and the ubiquitin-lifeactinGFP construct (Ubi-LifeActGFP, (Santa-Cruz Mateos *et al*, 2020)). To analyze F-actin distribution and dynamics in UAS-*α*PS1;UAS-*β*PS overexpressed FCs, tj-Gal4;Ubi-LifeActGFP females were crossed to UAS-*α*PS1;UAS-*β*PS males. All stocks and crosses were maintained at 25°C.

### Time-lapse image acquisition

For live imaging, 1–2 days old females were fattened on yeast at 25°C for 48–96 hours before dissection. Culture conditions and time-lapse microscopy were performed as described in (Prasad and Montell, 2007). Ovarioles were isolated from ovaries dissected in supplemented Schneider medium (GIBCO-BRL). Movies were acquired on a Leica SP5 MP-AOBS confocal microscope equipped with a 40× 1, 3 PL APO oil objective and Leica hybrid detectors (standard mode). Frames were taken every 30 s up to 1 hour. For each frame, eleven to twelve Z-stacks, with a 0.42 *μ*m interval, were captured to cover the entire basal surface of the cells.

### Image processing and data analysis

For quantification of basal actin dynamics over time, maximal projections of confocal stacks were created to account for egg chamber curvature. Integrated intensity of actin was quantified for manually selected regions using ImageJ software. Background value taken from cell-free region was subtracted from all data series. Data were subjected to Gaussian smoothening with s=3. A Moving average filter (second order) was applied to remove low amplitude noise. For quantification of integrin intensity along actin filaments, maximal projections of confocal stacks were produced to cover the entire basal actin organization. First, the Region of Interest (ROI) occupied by actin filaments was outlined by hand in each cell. ROIs were then transferred to the integrins fluorescence channel. Mean fluorescent intensity of integrins by pixel was calculated dividing integrated intensity by the total area occupied by actin filaments, using ImageJ software.

### Immunohistochemistry

Flies were grown at 25C and yeasted for 2 days at 25C before dissection. Ovaries were dissected from adult females at room temperature in Schneider’s medium to preserve cytoskeletal structures (Sigma Aldrich). Fixation was performed incubating egg chambers for 20 min with 4 % paraformaldehyde in PBS (ChemCruz). Samples were permeabilized using PBT (phosphate-buffered saline+0.1 % Tween 20). The following primary antibodies were used: chicken anti-GFP (1/500, Abcam), mouse anti-*β*PS (1/50, DHSB, Iowa). Fluorescence-conjugated antibodies used were Alexa Fluor 488 and Alexa Fluor 561 (Life Technologies). Samples were mounted in Vectashield (Vector Laboratories) and imaged on a Leica Stellaris equipped with a 40× 1, 3 HC PL APO CS2 oil objective.

## Acknowledgements

DGM acknowledges funding to the Spanish Ministerio de Econoíma y Competitividad (BFU2014-53299-P, RTI2018-096953-B-I00). PCF acknowledges funding to IFIMAC-UAM, through the ‘María de Maeztu’ Programme for Units of Excellence in R&D from the Spanish Ministerio de Economía y Competitividad (CEX2018-000805-M). ALI acknowledges funding to Comunidad de Madrid (PEJD-2018-PRE/BMD-7980). Institutional grants by Fundación Ramón Areces and Banco de Santander to the CBMSO are also acknowledged.

## Author Contributions

MH, AVE, PCF, and ALI performed research. PT designed research. MDMB performed research, designed research and provided funding, DGM performed research, designed research, provided funding and wrote the manuscript.

---

**Supplementary Figure 1.**
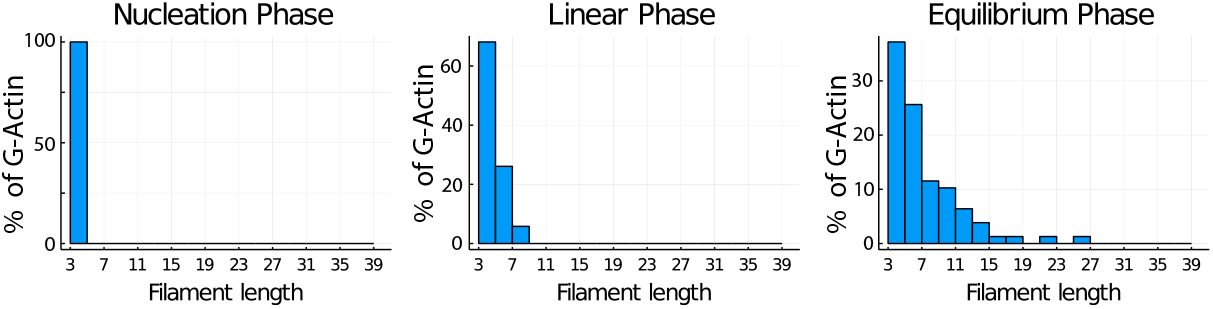
Percentage of G-Actin in filaments of different size for the three characteristic regimes.

**Supplementary Figure 2.**
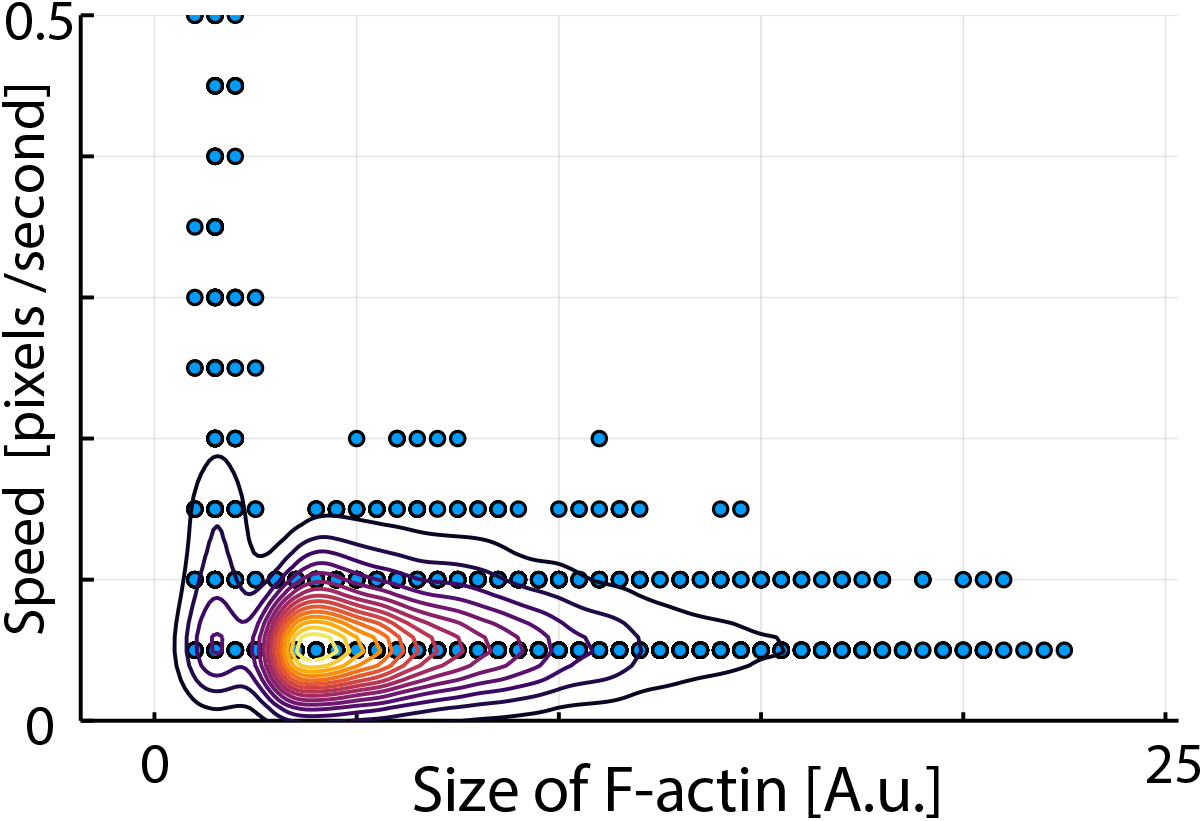
Dependence of speed of treadmiling with F-actin length. Plot of the dependence of the instantaneous speed on the filament size. Short filaments move faster than average. Long filaments move at the same speed in average.

**Supplementary Figure 3.**
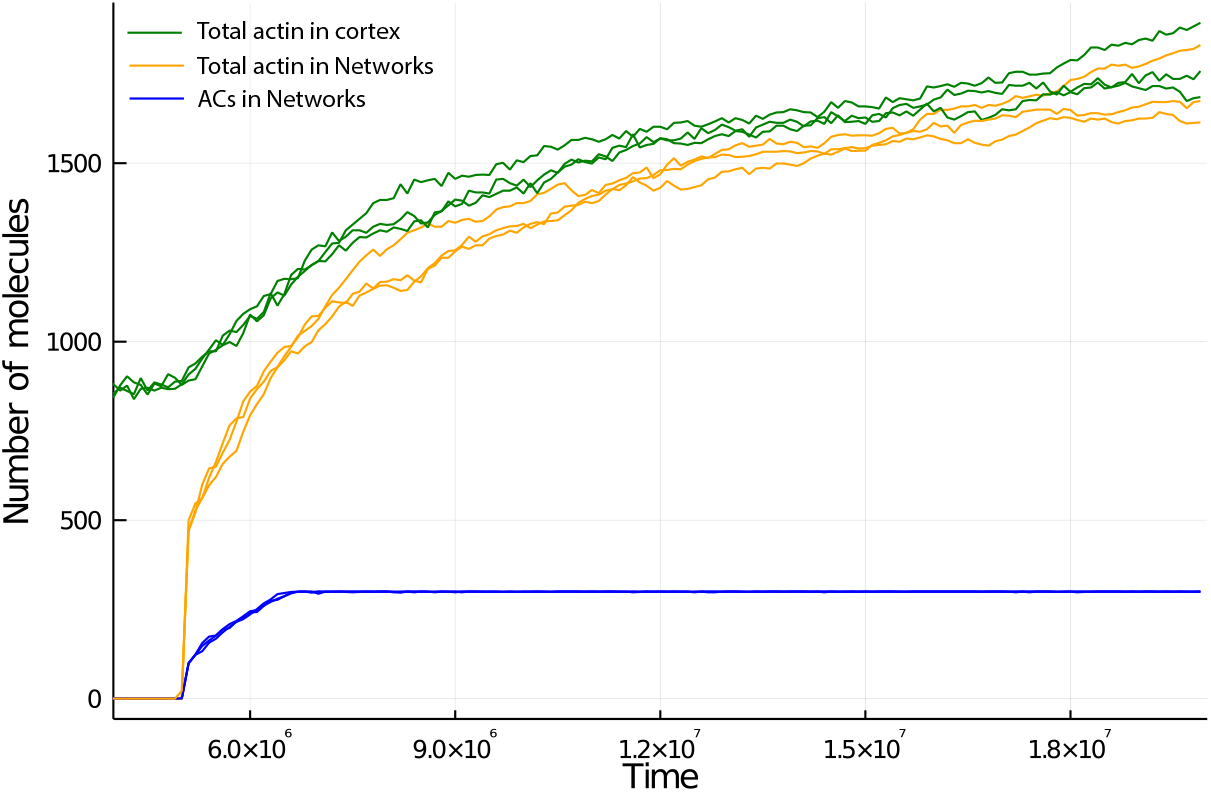
Number of G-Actin in the grid (green), G-actin as part of networks (orange) and ACs (blue) after incorporation of ACs into the system at t=5E6 iterations. The formation of networks is initial very fast, followed by a regime where incorporation of molecules into networks is gradually slowing down until equilibrium is reached.

**Supplementary Figure 4.**
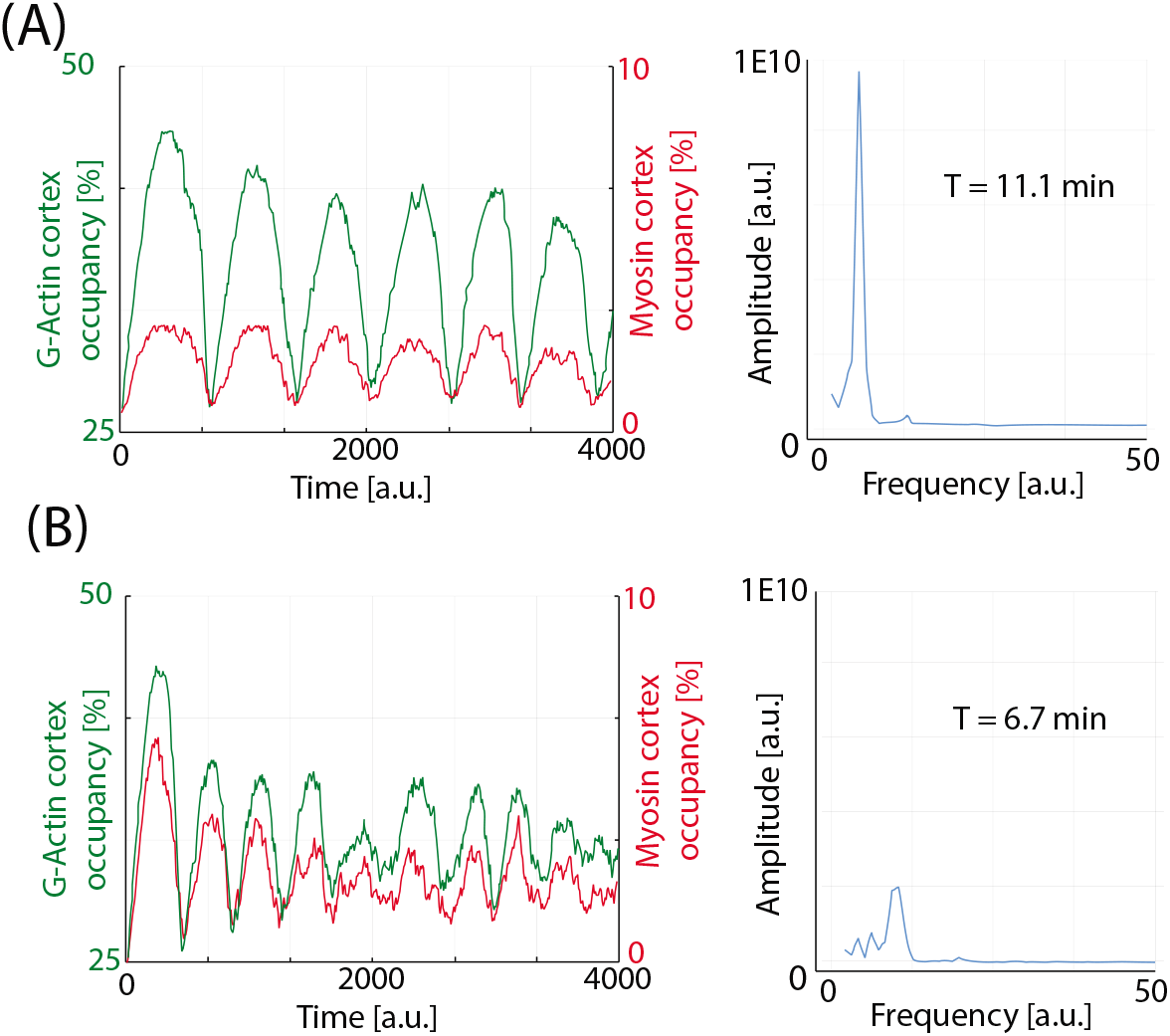
Oscillations in the number of G-actin in the cortex for conditions of (A) low and (B) high concentration of Myosin in the system. The corresponding Fourier transform for each oscillation is also shown.

**Supplementary Figure 5.**
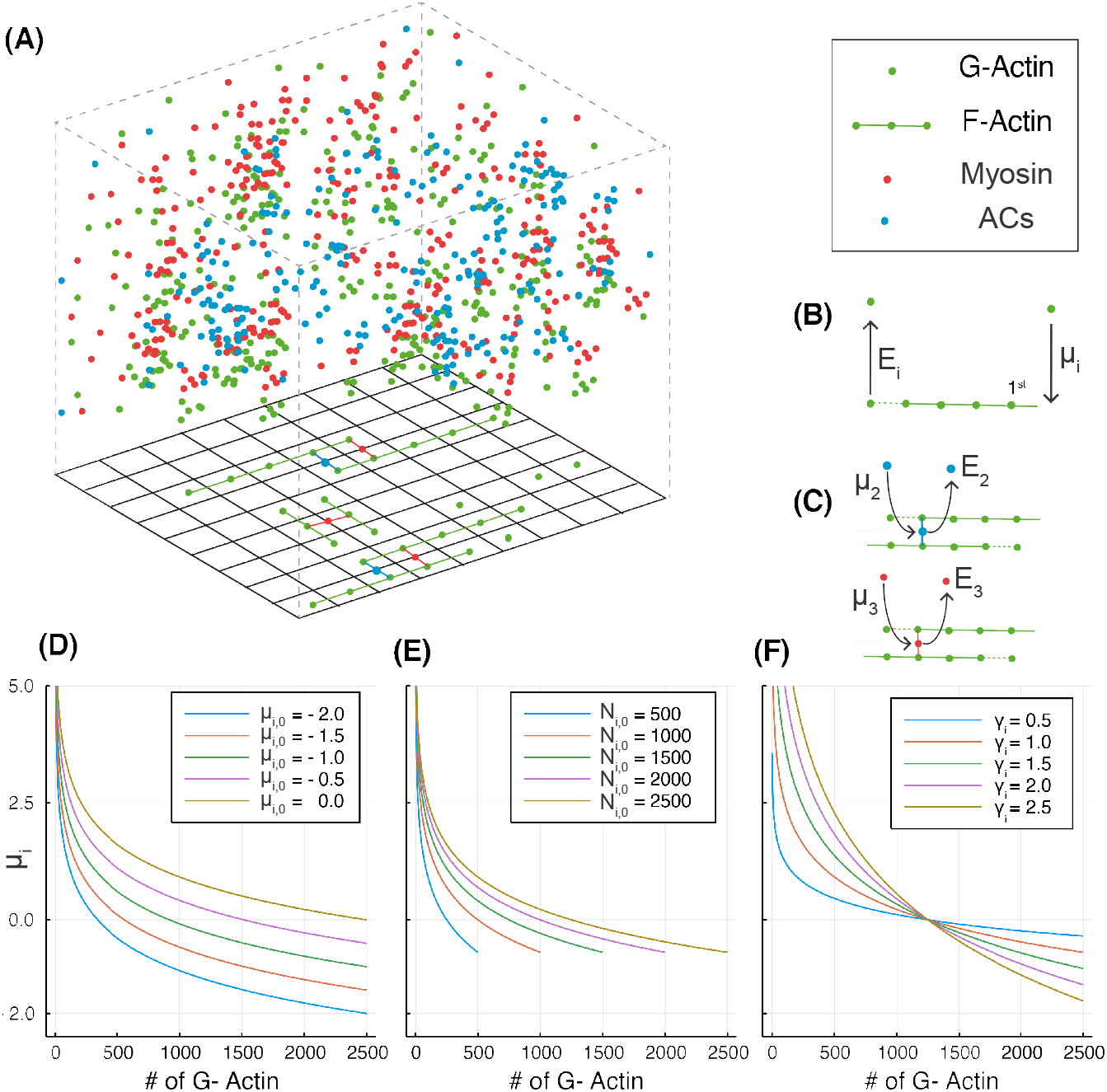
(A) Scheme of the framework. Molecules diffuse freely in a threedimensional space (cytoplasm) adjacent to a two-dimensional grid (inner plasma membrane) where molecules can attach. G-Actin (green) molecules in the grid interact and polymerize directionally to form F-Actin. ACs (blue) and Myosin (red) also interact with F-Actin to form networks of F-actin. (B) F-actin filament is formed by assembly at the barbed end (regulated by *μ*_1_) and disassembly at the pointed end (regulated by *E*_1_). (C) Linker formation of ACs and Myosin to F-Actin are regulated by *μ*_2_ and *μ*_3_, respectively. Release of ACs and Myosin is regulated by *E*_2_ and *E*_3_. (D-F) Shape of the potential function *μ*_1_ at a given time point for different values of (D) the reference potential *μ*_*i*,0_, (E) the total G-actin molecules in the system *N*_i,0_, and (F) the shape parameter *γ_i_*.

**Supplementary Figure 6.**
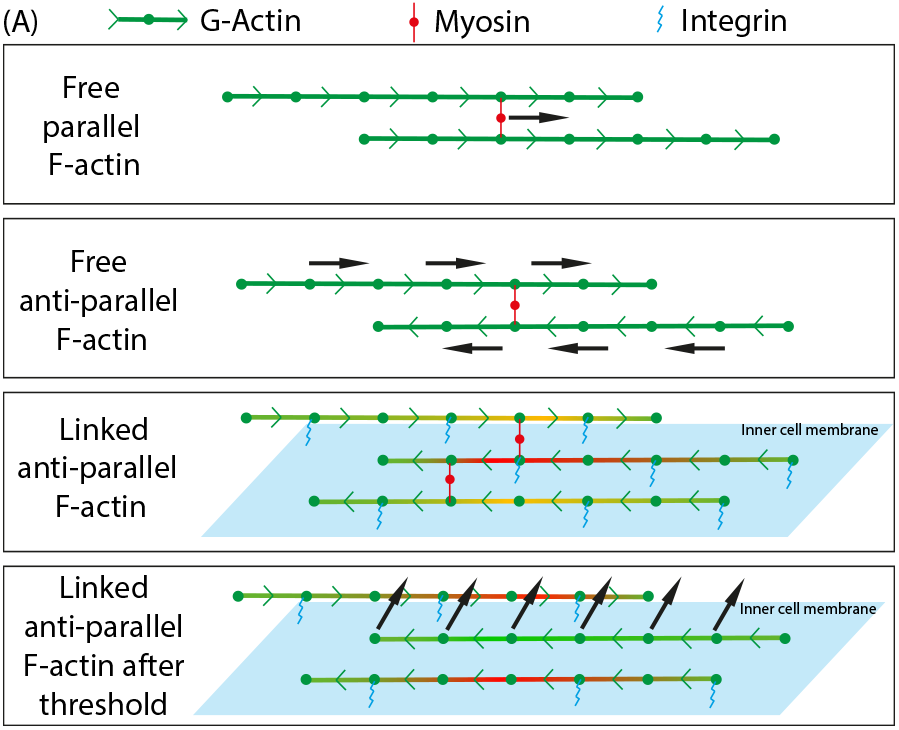
Scheme of the effect of Myosin over F-actin. (1), Myosin movement in parallel f-Actin. (2) F-actin sliding. (3) Tension building in the filaments. (4) Release from cortex after threshold tension is reached. After release, tension (illustrated in red) is redistributed to other F-actin in the network.

**Supplementary Movie 1.** Time-lapse movie of the system with F-actin forming and treadmilling in the grid.

**Supplementary Movie 2.** Time-lapse movie of the system with F-actin and ACs, where the formation of networks is taking place in the grid (G-actin labeled in green, ACs labeled in blue).

**Supplementary Movie 3.** Time-lapse movie of the system showing periodic assembly and disassembly of the actomyosin cortex (G-actin labeled in green, ACs labeled in red, Myosin labeled in red).

**Supplementary Movie 4.** Time-lapse movie of the D. Melanogaster basal follice cells stained with Lifeact-GFP in control conditions.

**Supplementary Movie 5.** Time-lapse movie of the D. Melanogaster basal follice cells stained with Lifeact-GFP in conditions of over-expression of the twe subuntis of teh integrin molecule.

**Table 1:** Values for the parameters used in the panels.

| Parameter Values for figures |  |  |  |  |  |  |  |  |  |  |  |  |
| --- | --- | --- | --- | --- | --- | --- | --- | --- | --- | --- | --- | --- |
| Figure | $\mu_{1,0}$ | $N_{1,0}$ | $\gamma_1$ | $E_1$ | $\mu_{2,0}$ | $N_{2,0}$ | $\gamma_2$ | $E_2$ | $\mu_{3,0}$ | $N_{3,0}$ | $\gamma_3$ | $E_3$ |
| Fig. 1 | -5 | 3000 | 2 | 4 | - | - | - | - | - | - | - | - |
| Fig. 2A | -5 | label | 2 | label | - | - | - | - | - | - | - | - |
| Fig. 2B-C | label | label | 2 | label | - | - | - | - | - | - | - | - |
| Fig. 3A | -4 | 2000 | 1 | 4 | - | - | - | - | - | - | - | - |
| Fig. 3B | -3 | 5000 | 1 | label | - | - | - | - | - | - | - | - |
| Fig. 3C | label | 5000 | 1 | label | - | - | - | - | - | - | - | - |
| Fig. 4A | -5 | 3000 | 2 | 3 | -5 | 300 | 2 | 3 | - | - | - | - |
| Fig. 4B | -2 | 3000 | 2 | 3 | -2 | label | 2 | label | - | - | - | - |
| Fig. 4C | -2 | label | 2 | 3 | -2 | label | 2 | 3 | - | - | - | - |
| Fig. 4D | -4 | 6000 | 2 | 3 | -4 | label | 2 | 3 | - | - | - | - |
| Fig. 5A | -3 | 5000 | 1 | 3 | -5 | 500 | 1 | 3 | -5 | 250 | 1 | 3 |
| Fig. 5B-D | -3 | 5000 | 1 | 1 | -5 | 3000 | 1 | 3 | -5 | 500 | 1 | 3 |
| Fig. 6A-B | -3 | 5000 | 1 | label | -5 | 3000 | 1 | 3 | -5 | 500 | 1 | 3 |
| Sup Fig 1 | -5 | 3000 | 2 | 4 | - | - | - | - | - | - | - | - |
| Sup Fig 2 | -3 | 2000 | 1 | 3 | - | - | - | - | - | - | - | - |
| Sup Fig 3 | -5 | 3000 | 2 | 3 | -2 | 300 | 2 | 3 | - | - | - | - |
| Sup Fig 4 | -3 | 2500 | 1 | 5 | -5 | 1250 | 1 | 3 | -5 | label | 1 | 3 |
| Sup Fig 5D | label | 2500 | 1 | - | - | - | - | - | - | - | - | - |
| Sup Fig 5E | 0 | label | 1 | - | - | - | - | - | - | - | - | - |
| Sup Fig 5F | 0 | 2500 | label | - | - | - | - | - | - | - | - | - |

