## Supplementary figures and images for "A coarse-grained approach to model the dynamics of the actomyosin cortex"

### SuppMovie1.gif

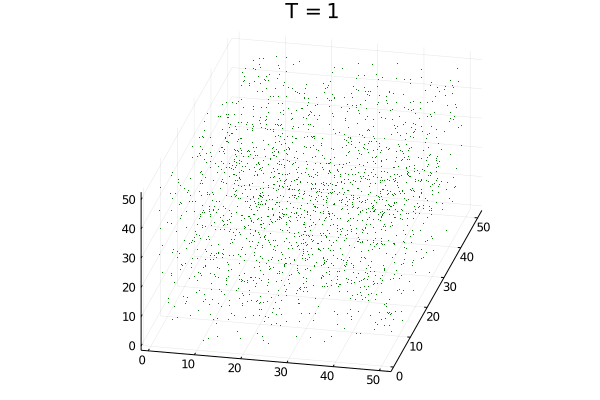

### SuppMovie2.gif

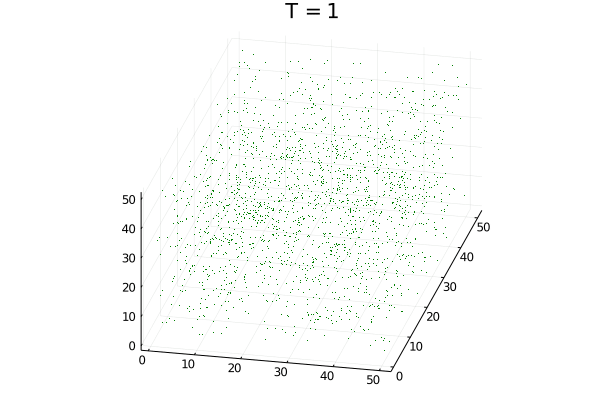

### SuppMovie3.gif

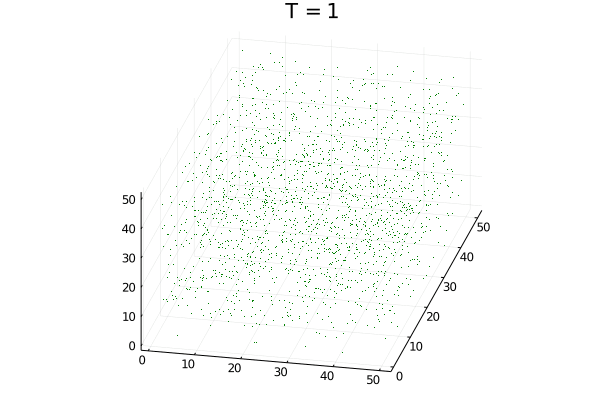
